# The climate benefits of yield increases in genetically engineered crops

**DOI:** 10.1101/2021.02.10.430488

**Authors:** Emma Kovak, Matin Qaim, Dan Blaustein-Rejto

## Abstract

The benefits of genetically engineered (GE) crops are systematically underestimated because previous studies did not incorporate the reduction in greenhouse gas (GHG) emissions associated with yield increases. We estimate this impact using the carbon opportunity cost of land use. Our results suggest that the GHG emissions reductions from the yield increases in GE crops are substantial and should be included in future analyses.

---

Previous studies quantified the economic benefits of GE crops, including increases in crop yields and profits, as well as the environmental and health benefits resulting from reduced pesticide use^1,2^. Lower GHG emissions from reduced tillage and savings in the use of tractor fuel were also considered^3^. However, reductions in GHG emissions associated with yield increases in GE crops were not quantified. As global demand for food and other agricultural products continues to grow, crop yield increases reduce the need to add new land into production, thus preventing additional CO_2_ emissions^4^ (Fig. 1). Land-use change accounts for almost half of all GHG emissions from agriculture^5^.

**Fig. 1.**
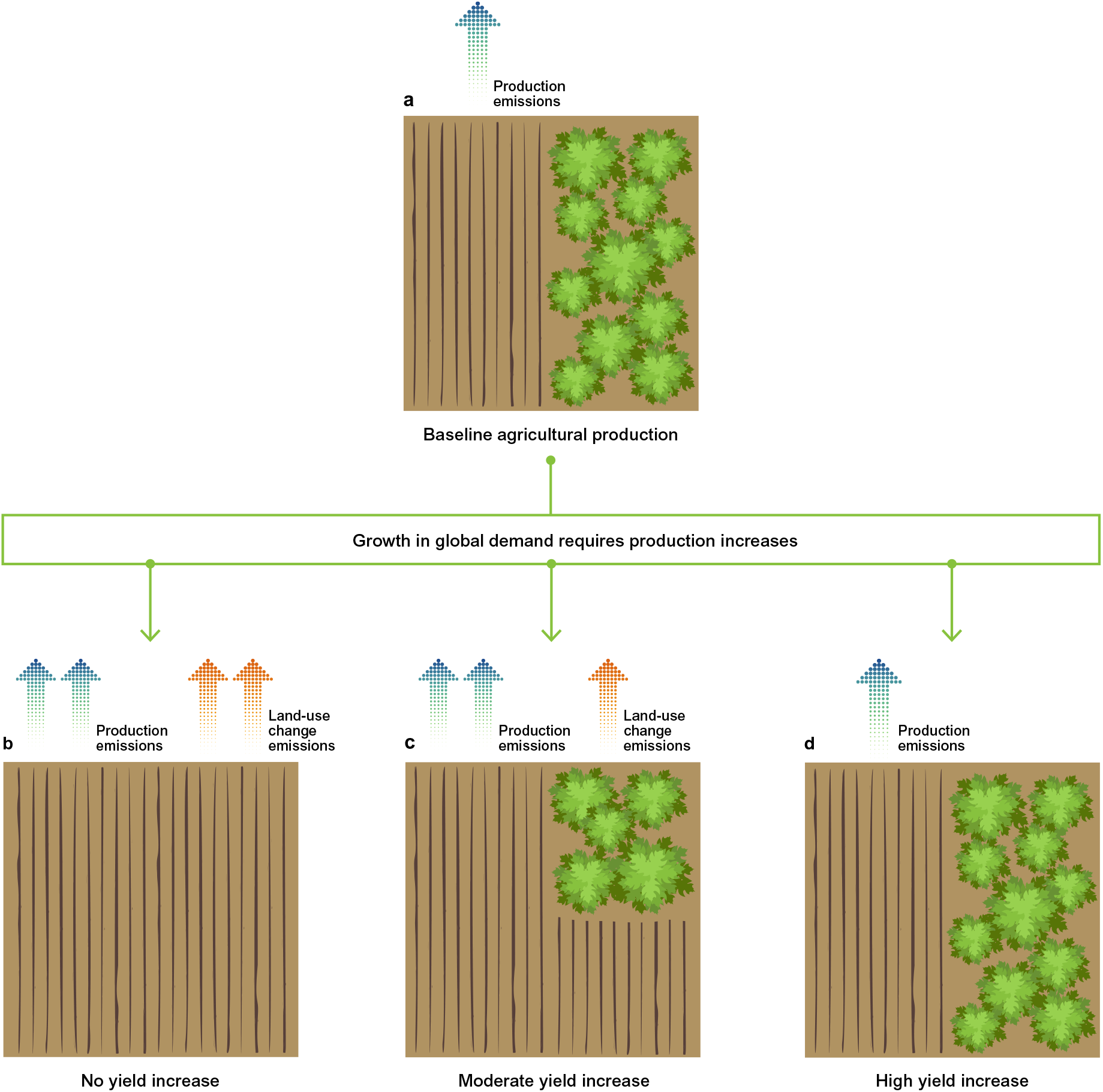
Increasing crop yields decrease the land conversion needed for agricultural production. This representation is simplified, as land use change and land sparing usually happen in a different location than original production. (a) Baseline agricultural production, which happens on a given amount of cropland with production-related GHG emissions. The rest of the land has native vegetation (forest, wetland, grassland, etc.). (b) Increased agricultural production without yield increase entails high conversion of natural land into cropland, with high land-use change emissions and increased production emissions. (c) Increased production with moderate yield increase results in moderate land-use change emissions and increased production emissions. (d) Increased production with high yield increase can help prevent both land-use change emissions and additional production emissions.

Here, we calculate the climate benefits of yield increases in GE crops, building on a method developed by Searchinger *et al*. (2018)^6^. We estimate to what extent GHG emissions could have been avoided if the European Union’s (EU) level of adoption of GE varieties of five major crops (maize, soybean, cotton, canola and sugarbeet) had been equal to that of the USA (Methods and Supplementary Tables 1 and 2).

While the climate benefits of GE crops can be calculated globally, we concentrate on the EU for two reasons. First, the EU has not yet widely adopted GE crops, mostly due to issues with public acceptance and related political hurdles^1^. This means that we can compare a hypothetical scenario with GE crop adoption to the status quo where hardly any GE crops are grown. Second, the EU is currently undergoing a reassessment of its GE policies (following Council Decision 2019/1904), and this analysis could help provide a more comprehensive picture of the likely effects of policy change.

Based on previous research we know that GE crops can increase yields, and we would expect this effect if the EU adopted GE crops. While various GE crop traits have been developed, the most widely adopted ones in different parts of the world are insect resistance (IR) and herbicide tolerance (HT)^1,7^. These traits help to reduce crop damage from insect pests and weeds, thus increasing effective yields. A global meta-analysis showed that the average yield advantages of GE crops are around 22%, with some differences between traits and geographical regions^8^. The average yield increase from GE adoption in temperate-zone industrialized countries is 9.7% and 6.5% for IR and HT, respectively^8^.

If EU crop yields increase with the hypothesized adoption of GE traits, we would expect this increased production in the EU to lead to a decrease in production and related land-use change elsewhere. In our analysis we consider two components of GHG emissions: the carbon opportunity cost (COC) of land use and production emissions (PEM). The COC is defined as the land’s opportunity to store carbon if it is not used for agriculture. This is influenced by the average carbon stocks in the native vegetation and in the soil, as well as the global average yield of each crop. Higher carbon stocks mean higher potential emissions from clearing land. Hence, shifting production towards places with yields above the global average – such as the EU – enables greater carbon storage on spared land elsewhere. PEM are calculated based on fertilizer and energy input use in production.

We find that growing GE crops in the EU could reduce GHG emissions by 33 million metric tons of CO_2_ equivalents per year (MtCO_2_e/yr). This is equivalent to 7.5% of total EU agricultural GHG emissions in 2017^9^. The avoided emissions per hectare are higher for maize than for the other crops (Fig. 2a), which is due to the fact that GE varieties with stacked IR and HT traits, and thus higher yields, are widely available for maize. Maize also accounts for the largest share (63%) of the total GE-related emission reductions in the EU (Fig. 2b, Supplementary Table 3), because maize is grown on larger areas than the other four crops considered. For all five crops, COC makes up a much larger proportion (>84%; Supplementary Table 3) of the total potential avoided GHG emissions than PEM, underlining the importance of considering COC when estimating the climate impacts of agricultural production and policy changes. For soybean, the smallest PEM reductions occur because average soybean PEMs in the EU are higher than the global average.

**Fig. 2.**
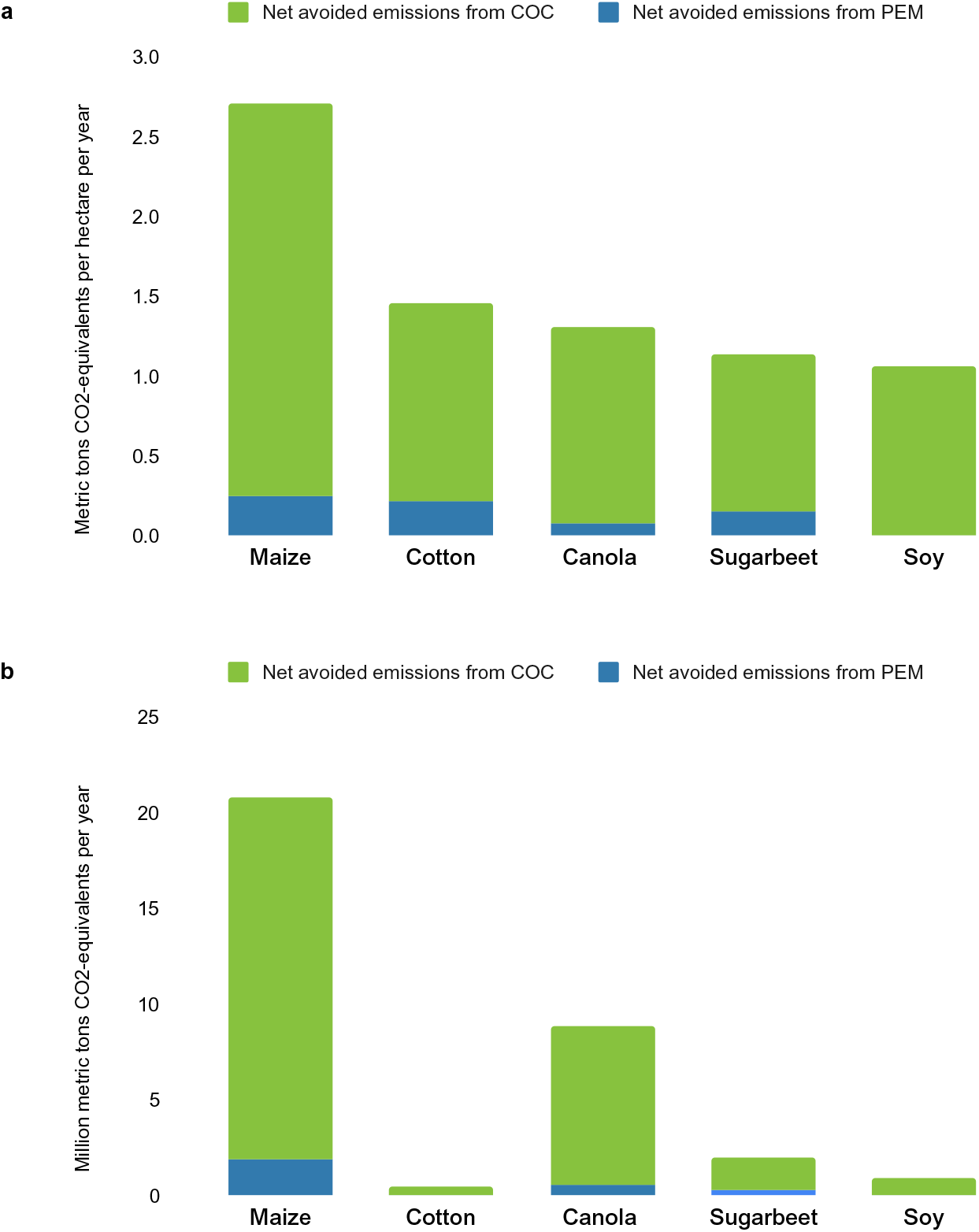
Potential avoided GHG emissions resulting from yield increases of GE crops in the EU. COC, carbon opportunity cost of land use. PEM, production emissions. (a) Estimates per hectare and year. (b) Estimates for total EU crop area per year.

Our analysis builds on a few assumptions that may lead to certain inaccuracies in the estimates. For instance, we assume that increased crop production in the EU leads to a proportional decrease in production elsewhere. While significant global land-sparing effects of crop yield increases are clearly established^4^, the magnitude can vary and is difficult to predict precisely in a particular situation^10,11^. Our assumption of a proportional link may lead to overestimation of avoided emissions. However, this is likely compensated for by two other assumptions that lead to underestimation. First, we assume that increasing GE crop adoption in the EU would have no effects on technology adoption in other countries, even though the quasi-ban of GE crops in the EU over the last 20 years has also discouraged their use elsewhere^1^. Hence, higher GE crop adoption in the EU would likely also lead to higher adoption elsewhere, including in Africa and Asia. Second, although we only consider mitigation potential in 2017 based on the adoption of IR and HT traits in five crops available at the time, future adoption of more GE crops and traits will likely increase emissions reductions, especially with more conducive EU policies.

Restrictive regulatory policies in the EU and elsewhere have limited the number of GE crops and traits developed globally^12^. Which crops have commercialized GE varieties is clearly impacted by the economic demands of countries that permit them. The fact that soybean, maize, and cotton are among the most widely-grown crops in the USA has certainly driven the creation of their GE varieties. By contrast, the most widely grown crops in the EU — wheat and barley — have no commercialized GE varieties, but this could change if European scientists had a market to work toward, particularly since Northwest Europe has levels of wheat crop losses due to pests and diseases that are above the global average^13^.

European researchers have argued repeatedly against the EU’s reticence to accept GE crops^14^. However, the EU may be headed in the opposite direction, as the new Farm-to-Fork Strategy under the European Green Deal aims to expand organic farming, which has lower yields and would be associated with significant increases in global GHG emissions through causing land-use change elsewhere^15^. Rather than offshoring environmental damage to other nations, as the European Green Deal does, the EU should increase agricultural productivity through embracing new crop technologies, thus contributing to global environmental benefits^16^.

There is reason to be hopeful. As crop biotechnology research continues, a wider variety of traits will become available, each with different yield impacts. Similar to IR and HT, GE crops tolerant to other stressors such as drought and heat will improve effective yields through reduced crop damage. Possibly more dramatic yield increases could come from GE traits that improve yield potential through enhanced plant growth and photosynthetic efficiency^17^. New gene editing technologies will likely further increase the diversity of desirable crop-trait combinations^12^. Larger yield increases in more crops would lead to larger GHG emission reductions. Hence, our estimate of 33 MtCO_2_e/yr is only a small proportion of the potential future benefits of GE crops for climate change mitigation.

## Methods

### Countries analyzed

We consider the EU with 28 member countries, before Brexit, as our analysis refers to 2017 as the base year.

### Crops analyzed

We include the five GE crops with the highest adoption rates in the USA (soybean, maize, cotton, canola, and sugarbeet), which includes the three most widely grown GE crops worldwide (soybean, maize, and cotton) plus two grown especially in the cooler temperate-zone climates of North America (canola and sugarbeet). In the EU, all soybean, cotton, canola and sugarbeet grown are non-GE varieties, and all but a tiny proportion of maize grown in the EU is non-GE varieties.

### Area cultivated and historical crop yields

For the total area cultivated for each crop and for historical crop yields, we use FAOSTAT data for the EU28 special grouping from 2017 (Supplementary Table 1).

### Adoption rates

In the four countries that plant the most GE crops worldwide — USA, Brazil, Argentina, and Canada — adoption rates for soybean, maize, cotton, canola, and sugarbeet range from 85–100%; we use adoption rates from the USA in 2017^7^ (Supplementary Table 1). As a robustness check, we re-calculate total avoided GHG emissions using lower adoption rates of 85% (the low end of all adoption rates in the top-four GE-growing countries worldwide) and 50% (a much lower adoption rate that might represent a country midway through increasing adoption). With 85% and 50% adoption for all five crops, total avoided GHG emissions are 30 and 17 MtCO_2_e/yr, respectively.

### Yield benefits of GE traits

We use data from a global meta-analysis of GE crop impacts to estimate yield increases^8^. Yield effects vary by geographic region. As we are interested in potential effects in the EU, we only consider the data from temperate-zone industrialized countries, not from developing countries where yield effects tend to be higher. We consider yield effects of IR and HT crops separately. For GE varieties with stacked IR and HT traits, we add the individual yield benefits. This assumption is reasonable, as insect pests and weeds cause separate yield damage, and farmers make decisions about insecticide and herbicide sprays independently. In their global analysis of GE crop impacts, Brookes and Barfoot (2020) also added the benefits of stacked IR and HT traits. As an additional robustness check, we calculate total avoided emissions using a lower value for the yield benefit of stacked traits — just the yield benefit from the IR trait, rather than adding the effects from IR and HT traits. This is only relevant for maize and cotton, as these two crops are the only ones for which stacked traits are available. Total avoided emissions in this alternative scenario are 25 MtCO_2_e/yr.

### Fertilizer application

We use 2011 data on nitrogen fertilizer application per hectare by country and crop from Zhang (2015) in addition to 2011 FAOSTAT data for the area of each crop harvested in individual EU countries. We summed the Zhang (2015) data for total nitrogen applied across all EU countries represented for each crop, then divided by the sum of the areas harvested from all EU countries that grew that crop (Supplementary Table 2). We incorporated this EU average value for nitrogen applied per hectare of each crop into the site-specific production emissions (PEM).

### Site-specific PEM

For the PEM (production emissions) component of calculating total avoided emissions, we used tabs 3.1 and 3.2 of the Searchinger (2018) Carbon Benefits Calculator. Users may enter site-specific values for all or a subset of fertilizer application, on-farm energy use, and energy use to produce fertilizer and pesticides. We entered site-specific values only for fertilizer application, and used default values for energy use because this component of total emissions only makes a small contribution compared to COC, so further site-specific values would make only a small difference in total emissions reductions. We used the default CO_2_e/N input, and we did not input values for rice methane. Site-specific changes in fuel use could also incorporate decreases due to reduced tilling and pesticide application, as calculated by Brookes and Barfoot (2020).

### Carbon Benefits Calculator

In order to calculate the total potential for avoided emissions from GE yield increases, we used tabs 1, 3.1, and 3.2 (the latter two as described above for PEM) of the Searchinger *et al*. (2018) Carbon Benefits Calculator to compute the avoided GHG emissions from the higher yields with GE varieties compared to the 2017 conventional yields per hectare of each crop. We used the default COC for each crop with the default 4% discount rate, fresh matter as the weight type for all variables, and did not enter inputs for livestock or bioenergy. Then we multiplied this difference by the total area of that crop harvested across the EU in 2017 adjusted for the assumed GE adoption rate.

## Acknowledgements

We thank Kenton de Kirby for providing valuable comments on the manuscript.

## Author contributions

DBR conceived the study, contributed to methods, and contributed to editing. EK contributed to methods, implemented the analysis, wrote the manuscript, and contributed to editing. MQ provided data and contributed to interpreting the results and to editing.

